# Meaningful Local Signaling in Sinoatrial Node Identified by Random Matrix Theory

**DOI:** 10.1101/2022.02.25.482007

**Authors:** Chloe F. Norris, Anna V Maltsev

## Abstract

The sinoatrial node (SAN) is the pacemaker of the heart. Recently calcium signals, believed to be crucially important in rhythm generation, have been imaged in intact SAN and shown to be heterogeneous in various regions of the SAN and shown to be heterogeneous in various regions of the SAN with a lot of analysis relying on visual inspection rather than mathematical tools. Here we apply methods of random matrix theory (RMT) developed for financial data and various biological data sets including *β*-cell collectives and EEGs to analyse correlations in SAN calcium signals using eigenvalues and eigenvectors of the correlation matrix. We use principal component analysis (PCA) to locate signalling modules corresponding to localization properties the eigenvectors corresponding to high eigenvalues. We find that the top eigenvector captures the common response of the SAN to action potential. In some cases, the eigenvector corresponding to the second highest eigenvalue yields a pacemaker region whose calcium signals predict the action potential. Furthermore, using new analytic methods, we study the relationship between covariance coefficients and distance, and find that even inside the central zone, there are non-trivial long range correlations, indicating intercellular interactions in most cases. Lastly, we perform an analysis of nearest-neighbor eigenvalue distances and find that it coincides with universal Wigner surmise under all available experimental conditions, while the number variance, which captures eigenvalue correlations, is sensitive to experimental conditions. Thus RMT application to SAN allows to remove noise and the global effects of the action potential and thereby isolate the local and meaningful correlations in calcium signalling.

## 1. Introduction

Calcium signalling is a universal property of excitable tissue, found for example in neurons in the brain, *β*-cell collectives in the pancreas, and in cardiac tissue, the subject of this study. The fundamental question regarding such tissue is how does the tissue ensure that action potentials are rhythmic, meaning close to periodic? The classical paradigm was that one cell or a small group of cells are the “pacemaker”, but with recent works this paradigm is starting to shift, in particular for neuronal [5], *β*-collective pancreatic [6], and cardiac tissue [4, 17, 3]. The new conception is that pacemaking is instead an emerging property of an interacting network of cells. In this paper we carry out a first mathematical analysis of intercellular calcium signalling in the sinoatrial node, based on recently obtained and published data of calcium imaging from the entire intact SAN [3] and on mathematical methods developed for *β*-cell collectives and financial assets [8, 12].

In the aforementioned recent experimental study [3], the authors introduced a novel imaging system with a high spatial and temporal resolution to record intracellular calcium dynamics and action potentials within an entire intact SAN. They established that an action potential (AP) induced calcium transients closely follow corresponding action potential onset, which means that a study of whole SAN intracellular calcium signals is sufficient to make conclusions about cardiac pacemaking. They immunolabelled HCN4 and highly conductive intercellular channels CX43, and established the existence of the central zone where cells express only HCN4 and the “railroad tracks” zone where cells express only CX43. Furthermore, using case studies of regions in the central zone identified by visual inspection, the authors found that the central zone features local calcium signals which are heterogeneous, and deduced that this heterogeneity is important for pacemaking. However, calcium signalling in the SAN central zone is very noisy and objective algorithmic identification of meaningful signaling domains and their further properties has not been developed.

The mathematical analysis of intercellular signalling is reasonably well-developed in *β*-collectives and their methodology could pave the way for an integrated mechanistic understanding of how the interacting SAN cells produce the working SAN. Calcium imaging video data from in situ pancreatic *β*-cells was first obtained almost twenty years ago in [14] using a tissue-slice technique that is classic in neuronal tissue. Since then in a series of papers, various network and statistical properties of *β*-cell collectives have been established [15, 7, 8]. In particular, using an analysis of calcium signal correlations, it has been shown that their network topology is small-world, which naturally arises in a multitude of settings ranging from social networks to the nervous system, demonstrated for example on C. *elegans* [16]. A key feature of small-world networks are highly connected “hubs,” which in *β*-collectives turn out to be crucial for pancreatic responses to glucose [7].

Another approach taken in analyzing calcium imaging data in *β*-collectives comes from random matrix theory [8] and its applications to finance [12]. This is the approach we take here. In this paper, we use random matrix theory and related tools such as PCA to investigate the correlation structure of the calcium signals in SAN. By studying the eigenvalues and eigenvectors of the covariance matrix, we are able to separate the eigenvalues arising from noise (those that follow RMT predictions), those that reflect long range correlations caused by propagating action potentials (top eigenvalue), and those reflecting local calcium signalling. While most eigenvalues follow RMT predictions, it is the few that deviate that carry meaningful information about local calcium signals. We also demonstrate that the eigenvalue spacing for the eigenvalues predicted by RMT follows the Wigner surmise, which is a universal distribution for eigenvalue spacing that arises in theoretical mathematics, as well as in numerous real-world examples including spacings of energy levels in heavy atoms, bus timings in Cuernavaca, Mexico [9], correlations in EEG data [13] as well as the two systems that inspired this work: financial markets and *β*-collectives. The fact that the eigenvalue distances follow the Wigner surmise confirms that the RMT-predicted eigenvalues correspond to system noise, and this noise has universal properties i.e. it is not special or unusual. We also examine the number variance and find that it is far more sensitive to experimental conditions, in line with prior work on human EEGs [13] and pancreatic *β*-cells [8].

By analysing the eigenvalues and eigenvectors that do not follow the random matrix theory predictions, we establish non-trivial properties of correlations of calcium signals. Using the localized eigenvectors associated to the high but not the highest eigenvalues, we can isolate correlated modules of points corresponding to signalling cellular modules. This approach is a powerful tool in data analytics, as it underlies PCA, which has been used successfully to identify various biological structures []. In one of the analyzed videos, we find that the calcium signal from the module that corresponds to the second-largest eigenvector just precedes the global action potential, thus suggesting that this may be either the pacemaker or the signal integrator module of the SAN.

Using RMT, we are able to remove the global effects of the action potential, to isolate the correlations in calcium signalling which are local. We investigate the spatiotemporal length of the correlations, obtaining, of course, that they are strongest at intra-cellular distances. However, we demonstrate the presence of weak correlations at distances that exceed cell width, suggesting intercellular communication despite the absence of gap junctions. This methodology of studying distance-dependent correlations is new to this paper and can be useful for other settings, such as in particular *β*-collectives.

This paper offers a first mathematical approach to analyzing SAN calcium imaging data. This is important because these data analysis methods are reproducible and more objective than the case studies used to study calcium signals in SAN previously.

## 2. Methods

### 2.1. Random Matrix Theory

The behavior of covariance matrices of purely random data is well-understood. In this paper, we rely on prior work in random matrix theory. The distribution of eigenvalues and various properties of eigenvectors are well-known when the matrix has independent identically distributed (iid) entries. Let *X* be a *T* × *S* matrix with independent real entries *x_ij_* with mean 0 and variance 1 (here mnemonically *T* stands for time and *S* stands for space). The sample covariance matrix is defined as *M* = *X^t^X*. Looking at the individual entries of the covariance matrix we see that 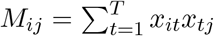, which tends to the Normal distribution with mean 0 and variance 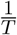 by the Central Limit Theorem.

Let *q* = *S/T* and 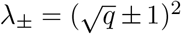. Then in the limit where *q* is constant and *T* → ∞, the density of eigenvalues *ρ_MP_* tends to the Marchenko-Pastur distribution supported on (λ_−_, λ_+_), namely for *x* ∈ (λ_−_, λ_+_)

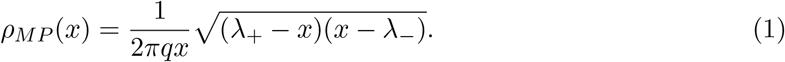

and *ρ*(*x*) = 0 for *x* outside (λ_−_, λ_+_) [10]. The eigenvectors of **C** are delocalized, meaning that none of their components are too big, and their components are Normally distributed. To study delocalization quantitatively, one introduces the inverse participation ratio (IPR) of a vector **v** of length S and norm 1, i.e. 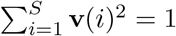, as

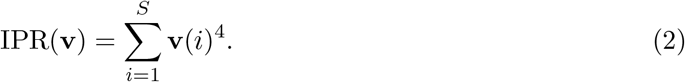

We notice here that the IPR can range between 1/*S* for vectors which are completely delocalized with each component equalling 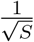 and 1 for vectors which are completely localized with one component equal to 1 and the rest equal to 0.

Lastly, the distances between nearest eigenvalues are also well-understood. They were studied by Wigner in a random matrix model where eigenvalues corresponded to energy levels in a heavy nucleus and the random matrix theory prediction described the repulsion between them (i.e. they two quantized energy levels cannot be too close to each other). The probability density *P* for the distances between nearest eigenvalues is universal for many ensembles, once the eigenvalue density is accounted for. This normalization for the eigenvalue density is called spectral unfolding, see Section 3.1.2 for details of unfolding. The density *P* is then given by the “Wigner surmise” equation see e.g. Section 1.5 of [11]

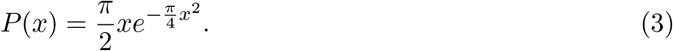

To probe the finer structure of eigenvalue correlations, one introduces the number variance Σ^2^(*L*). Letting ξ be an unfolded eigenvalue and 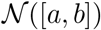 be the number of eigenvalues in the interval [*a*, *b*], the number variance is given by

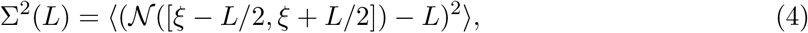

where in random matrix theory 〈·〉 denotes the ensemble average, while in analyzing data, the average is taken over unfolded eigenvalues ξ_*i*_ in the middle of the eigenvalue distribution. The random matrix theory prediction for the number variance is

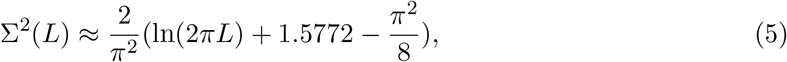

see e.g. Section 16.1 of [11].

We construct a matrix of covariances at various points from the calcium imaging data eigenvalues. Then we compare the properties of this matrix to the random matrix theory predictions. The features that fall within the random matrix theory predictions can thus assumed to be due to noise, while those that deviate can be interpreted as carrying meaningful information about spatial correlations of calcium signals, yielding insights into features of signal propagation.

### 2.2. Data sets and sampling

In this paper, we rely on calcium imaging data published in [3]. The intact SAN was flattened and a novel imaging system was used to assess calcium signaling. A membrane-permeable calcium indicator Fluo-4 was used in all videos, except Video 4 (also corresponds to Video 4 in this paper) where calcium signals specific to HCN4+ cells were recorded using SANC from mice (pCAGGS-GCaMP8) with HCN4-targeted expression of a calcium probe. Video 3 in this paper contains data about human SAN and corresponds to Video 5 in [3], while all others are from mouse. For details of SAN preparation and microscopy, please see Methods in [3].

The cells in the SAN divide into two disjoint types. One type of cell has the gap junction CX43 and no HCN4 gene, while the other cell type has the HCN4 gene and no gap junction CX43 [3]. We identify the central SAN that has no gap junctions by inspection, and take a convenient subset of it as our region of interest (ROI). Figure 1A shows the ROI that we work with for Video 1 (same as Video 1 in [3]) and Video 2 which is taken from the pptx supplemental material in [3].

**Figure 1.**
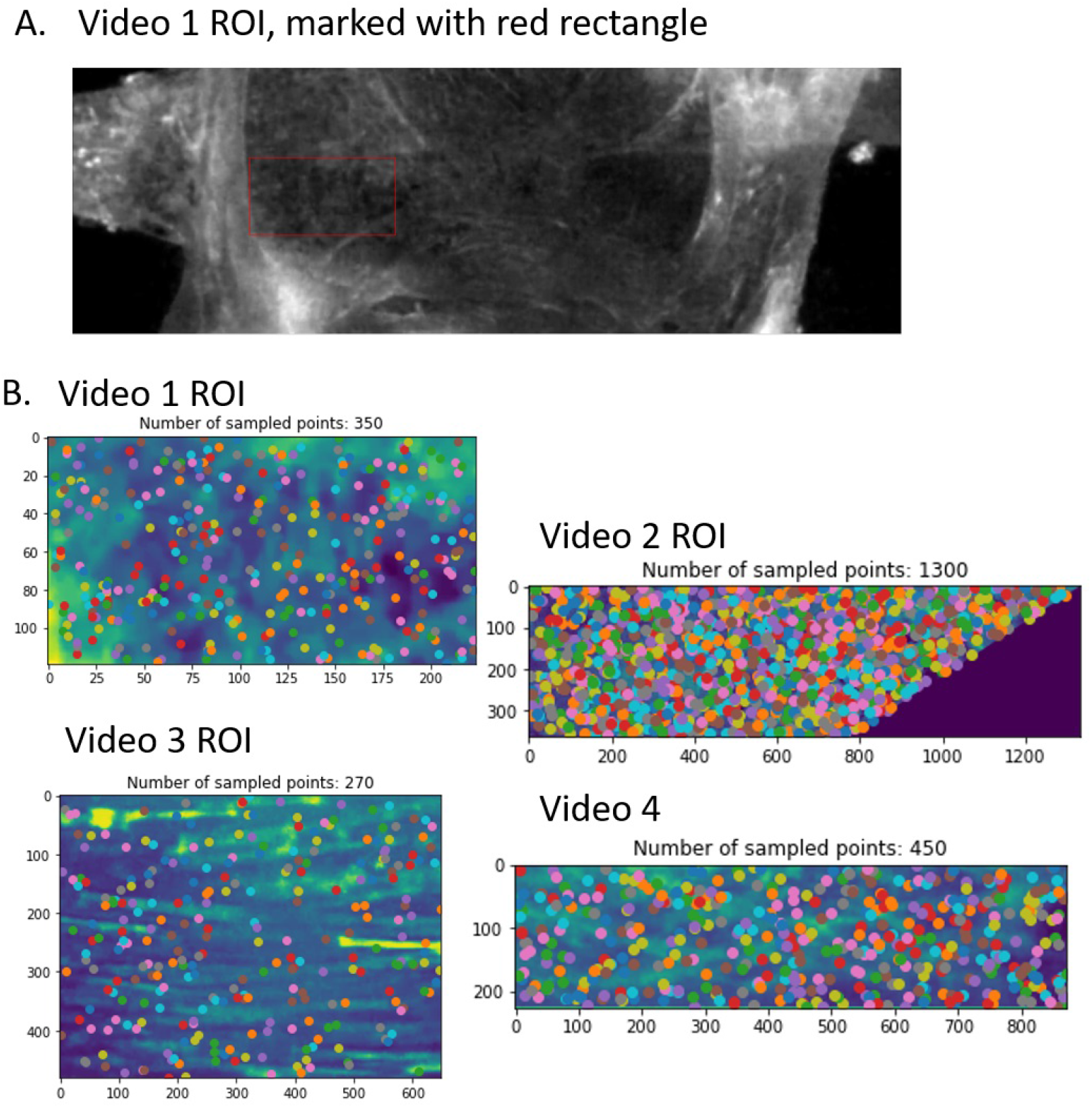
A: Choice of ROI from calcium imaging data of the SAN; B: Choice of a uniform random sample from Videos 1-4

In this paper, we study several data sets. As the videos are quite short, i.e. the number of time frames is much smaller than the number of pixels, and for computational efficiency, we must choose only a subset of the pixels, with each pixel corresponding to a time series. To avoid subjectivity, we do not identify cells in the SAN. To understand spatial properties of the calcium signals, we need to access the relationship between distance between pixels and covariance of the corresponding time series. Thus, we follow [8] in sampling the pixels uniformly at random so as to have a range of various distances available (picture of sampled points for each video can be found in Fig 1B). A grid sampling in this case would not be appropriate as pixels in a grid have a minimum distance. We let *S* be the sample size. For our analysis of signaling cell modules, we also compare our findings from the uniformly random sample with a grid sample.

We notice that a key assumption in the random matrix theory predictions is that the random variables have mean 0 and variance 1. Thus, upon obtaining our sample of time series, we normalize and demean it as is standard in financial literature, see e.g. equation (2) in [12]. Let time series *V_i_* be a time series in our sample with the index *i* running from 1 to the sample size, where *V_i_* is a function of *t* with *V_i_*(*t*) the magnitude of the *i*th signal at time *t*. Then we define and work with *υ_i_* as follows

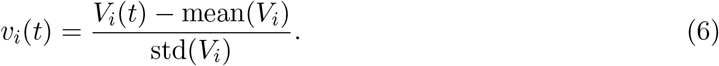

This procedure yields time series *υ_i_* with mean 0 and variance 1, and thus the features of covariances of these time series can be meaningfully compared to the random matrix theory predictions. The covariances of normalized and demeaned time series are the same as the correlations of the original time series.

In some cases, we also work with a time-incremented time series. This is analogous to working with returns rather than prices in financial mathematics, which decreases the impact of correlations within a time series on the analysis. We define the series 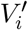 by 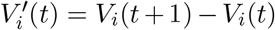. Then we work with 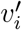, where 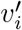 is 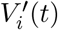 demeaned and normalized using (6). It is important that the time incrementing happens first, and the demeaning and normalization second, so that the resulting variables are of mean 0 and variance 1. Arranging the column vectors *υ_i_* in some arbitrary order to form the *T* × *S* matrix *V* we study the matrices

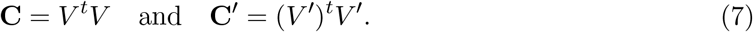

Upon examining the eigenvalues and eigenvectors of **C** and **C′** we find that the largest eigenvalue deviates from the random matrix theory predictions by a vast amount and the corresponding eigenvector is delocalized. We demonstrate that this corresponds to system response to a common stimulus, in this case the action potential propagating almost instantly via CX43. As we are interested in local rather than global signals we want remove the system response. We thus define another matrix of study in the following way, similar to [8]. Letting *u*(λ_*max*_) be the eigenvector corresponding to the largest eigenvalue λ_*max*_ with *u_i_*(λ_*max*_) referring to the components, we define

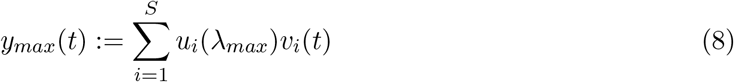

the projection of all the time signals in a particular pixels onto the top eigenvector. Then using linear regression on each time series *υ_i_* we obtain

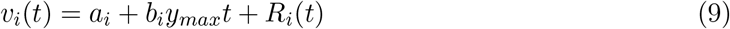

where *a_i_* and *b_i_* are coefficients of linear regression and *R_i_*(*t*) is the residual. The *T* × *S* residual matrix *R* given by *R_it_* = *R_i_*(*t*) forms the new matrix for analysis, where all the original information remains intact except the influence of the top eigenvector is removed. Thus let

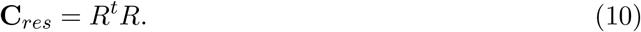

Lastly, in all our results we compare the properties of the data set with a control. As control we take the same matrix *V* and take a random permutation of the entries. We call this control matrix *V_perm_*, and define

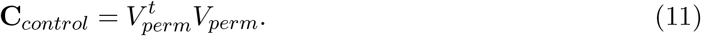

This procedure destroys all correlations within the data, while keeping the properties of the individual random variables intact. Comparing spectral properties of **C** to ones obtained from *V_perm_* allows us to rule out the possibility that any anomalies are due to properties of the distribution of individual entries, such as heavy tails.

## 3. Results

### 3.1. Eigenvalues and Eigenvectors

#### 3.1.1. Eigenvalue distributions

Looking at the histograms of eigenvalues (Fig. 2), we see perfect agreement of our control with the random matrix theory predictions with eigenvalues located in the predicted interval [λ_−_, λ_+_]. The histogram of the eigenvalues of **C** is also in agreement, but with a significant number of eigenvalues (≈ 3 – 6%) above [λ_−_, λ_+_]. We also notice a much better agreement with random matrix theory predictions for the time incremented matrix **C′**, demonstrating that time incrementing destroys some of the correlations within the data. We also run basic checks to ensure that these properties do not depend overly much on the random sample of pixels that we have checked the basic properties of the eigenvalues and eigenvectors in 500 randomly chosen samples for Videos 1 and 4, see Fig 2B. The box-and-whisker plot in Fig 2B shows the distribution of the top eigenvalue which is proportionately close to the mean. The histogram in Fig 2B shows that the number of eigenvalues outside the random matrix prediction range [λ_−_, λ_+_] is also reasonably stable.

**Figure 2.**
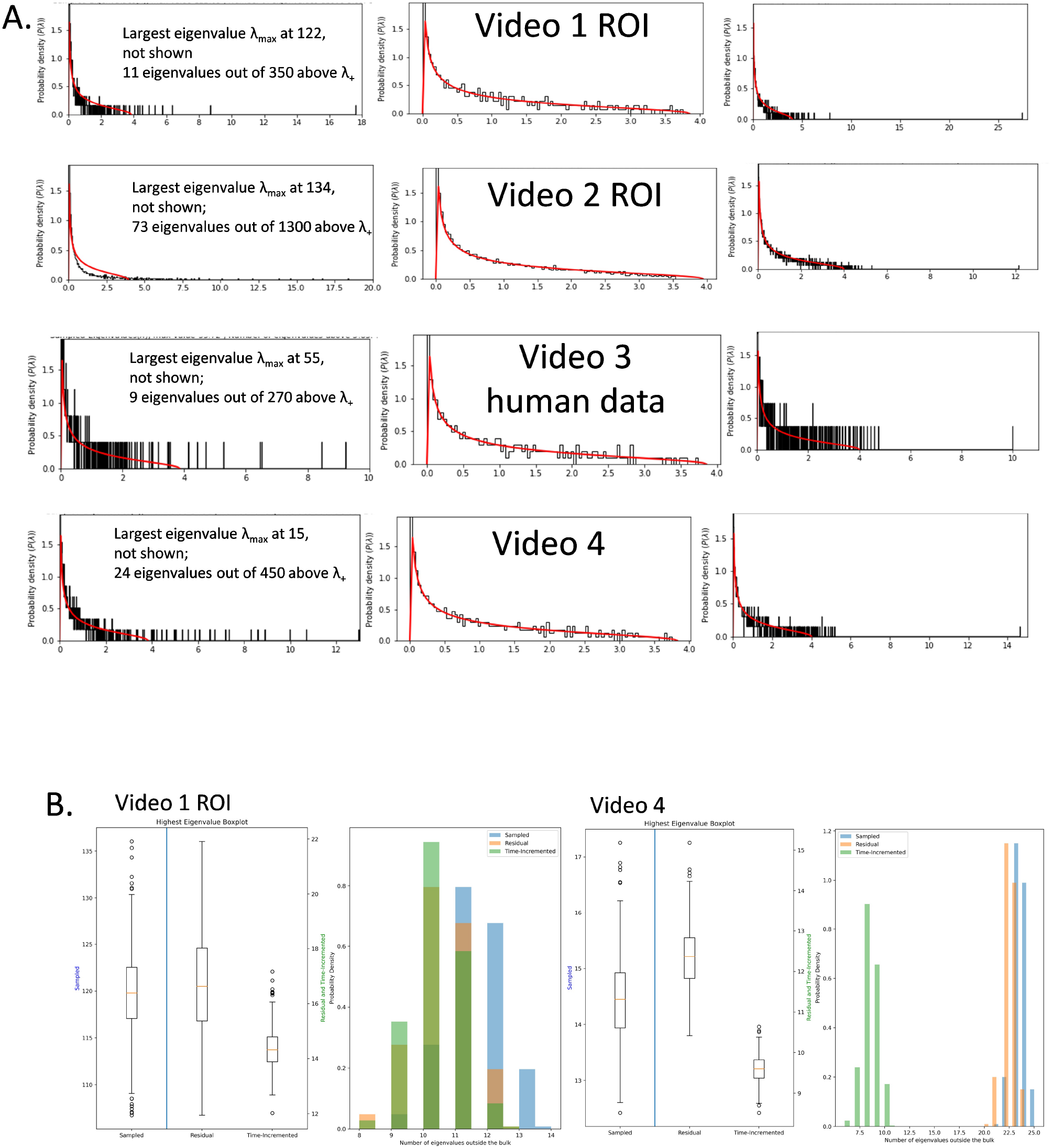
A: Eigenvalues distributions of the correlation matrix for Videos 1-4. Left column are eigenvalues of **C**, middle is of **C**_*control*_, right is of **C′**; B: stability analysis for various eigenvalue properties with 500 different random samples chosen for Video 1 and Video 4. On the left is a box plot for the top eigenvalue and on the right is for the number of eigenvalues outside of [λ_−_, λ_+_] for **C**, **C**_*res*_, and **C′**.

#### 3.1.2. Wigner Surmise

We study the distribution of distances between the eigenvalues in Fig 3A. To obtain a universal distribution for the distances between eigenvalues, we have to account for the eigenvalue density, called unfolding of the spectrum. We work with the eigenvalues of the time incremented matrix **C′**, since they conform to the Marchenko-Pastur distribution better than the eigenvalues of **C**, making the unfolding easier. We unfold the spectrum by using the Marchenko-Pastur law (1) as our limiting density, and we implement a classic procedure whereby we map each eigenvalue *E_j_* to a new variable 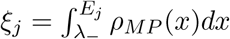. In this analysis, we neglect the eigenvalues *E_j_* < λ_−_ and *E_j_* > λ_+_. The resulting histogram of distances between nearest eigenvalues, avoiding largest 20% and smallest 20% of eigenvalues, is shown in Fig 3A in blue. We perform the same procedure for the eigenvalues of the control matrix **C**_*control*_, and it is plotted in Fig 3A in pink. We observe a close match to the Wigner surmise distribution in all analyzed videos, with a somewhat better match for the control case.

**Figure 3.**
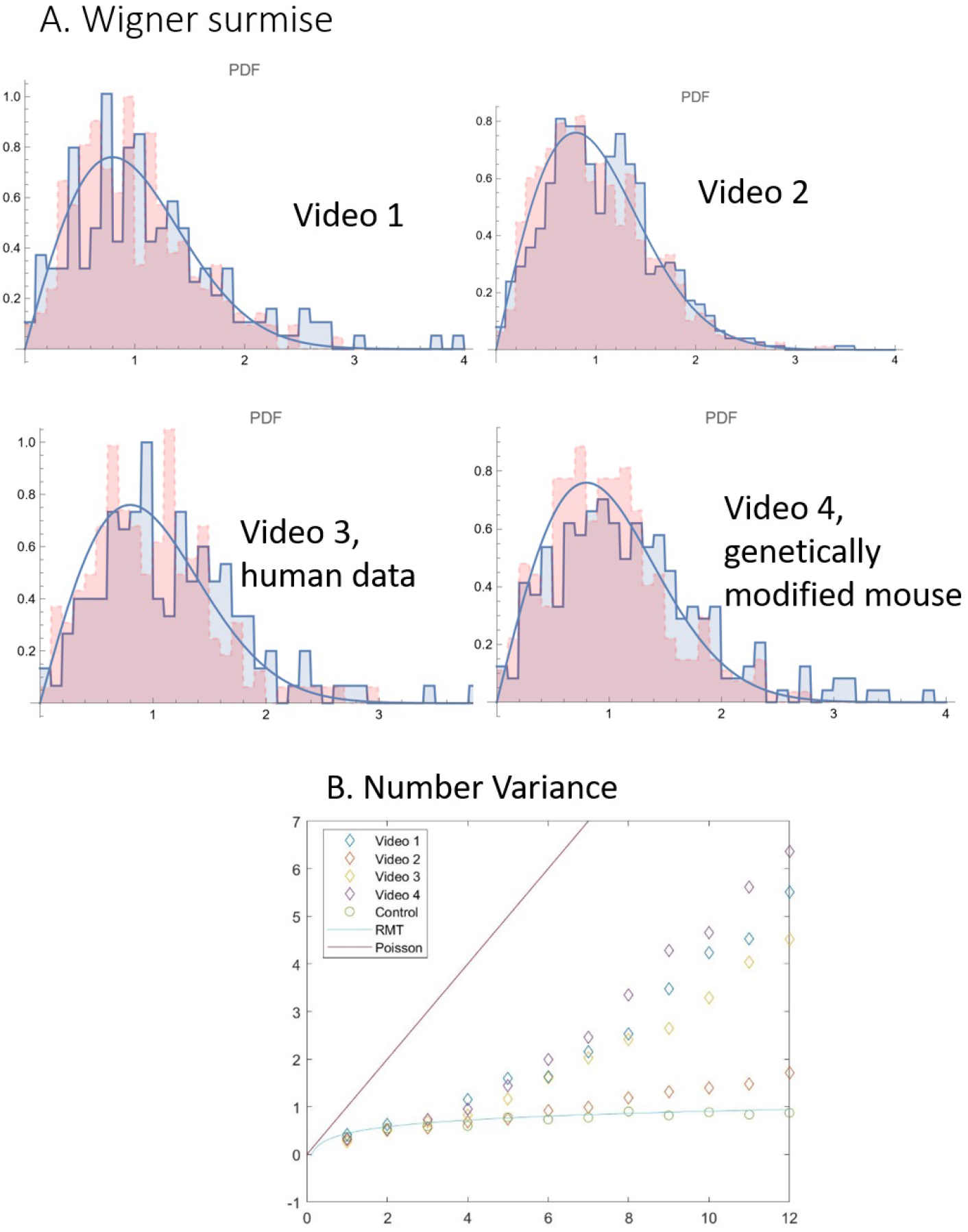
A: Distribution of distances between nearest unfolded eigenvalues from Videos 1-4, blue for unfolded data, pink for shuffled control, red for RMT prediction; B: the number variance for Videos 1-4, with red line for Poisson Process, blue line for RMT predictions, and circles for shuffled control

We then proceed to compute the number variance Σ^2^(*L*), which captures higher order correlations between the eigenvalues (Fig 3B). We use the same unfolding procedure, and only average over the middle 60% of eigenvalues to avoid edge effects. We see significant deviations from both the RMT prediction as well as from the Poisson process, thus the eigenvalues are correlated but not as much as eigenvalues of random matrices. This is in line with prior research on EEGs and *β*-cell collectives [8, 13].

#### 3.1.3. Inverse Participation Ratio and eigenvector localization

Next we examine the IPRs (Fig. 4). In all the diagrams we see that the control IPRs (in orange) are low, thus the eigenvectors are delocalized and follow the random matrix theory predictions. The IPRs are higher for eigenvalues larger than λ_+_ except at λ_*max*_. The higher localization at high eigenvalues indicates high correlations of particular pixels. The pixels where the eigenvectors of **C**, associated to eigenvalues above λ_*max*_, are localized thus indicate places where calcium signals move together. Lastly, we see that the eigenvector associated to the highest eigenvalue is fully delocalized. This indicates a common response of the cell system to a common stimulus, in this case the action potential transient.

**Figure 4.**
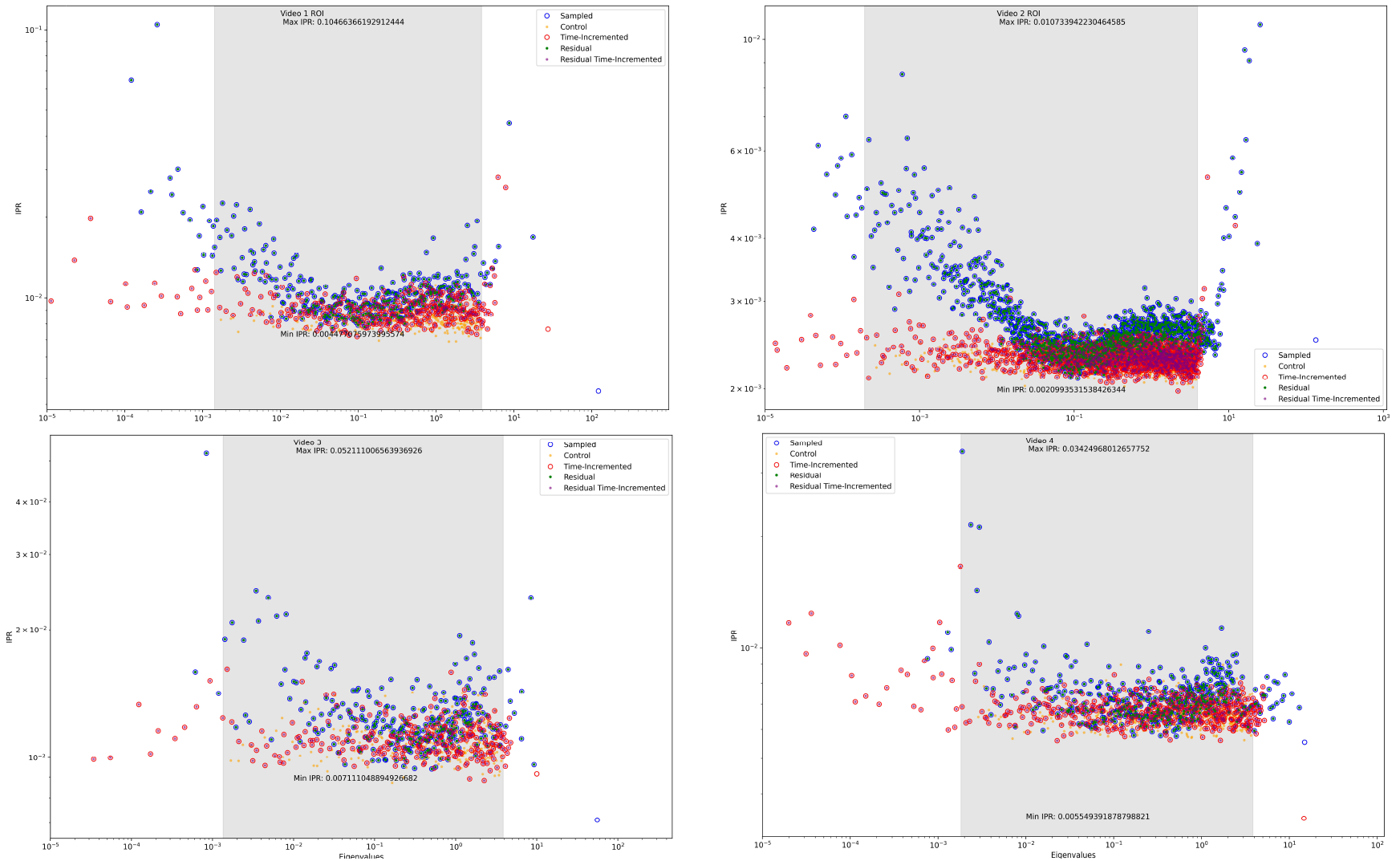
The IPR for Videos 1-4, calculated separately for the control, sampled, time-incremented, residual, and residual time-incremented data. Time-incremented residual data was calculated by adjusting around the lowest IPR from the eigenvalues outside of RMT predictions. The *x* axis and *y* axis are log scaled. IPR values are on the *y* axis, and eigenvalues are on the *x* axis. The areas of the graph that are shaded in grey is the area bound by RMT prediction.

In PCA, the eigenvectors that correspond to high eigenvalues of the data correlation matrix are called the principal components. Analogously to [12], we identify the locations where the principal components are localized. In the study of cross correlations for financial returns, the places of principal component localization corresponded to various industries: such as companies with high market capitalization, semiconductors-computers, gold, Latin-American firms, etc. In our case, these will correspond to cell signalling modules. Since the eigenvectors are orthogonal and localized, these modules will be non-intersecting. In Fig 5, we see the eigenvector components plotted in an arbitrary order, with a 2 standard deviation threshold. The components which exceed the threshold are places of eigenvector localization and when mapped back to the videos will signify signalling modules. We identify them in Video 1, Video 2, Video 3, and Video 4. Since we have taken a random sample of pixels, we map back five different samples to make sure that these identified signalling modules are not an artifact of a particular random sample. We see that the resulting modules are quite close (Video 5); they are also quite close to those identified by a grid sample, see Video 6.

**Figure 5.**
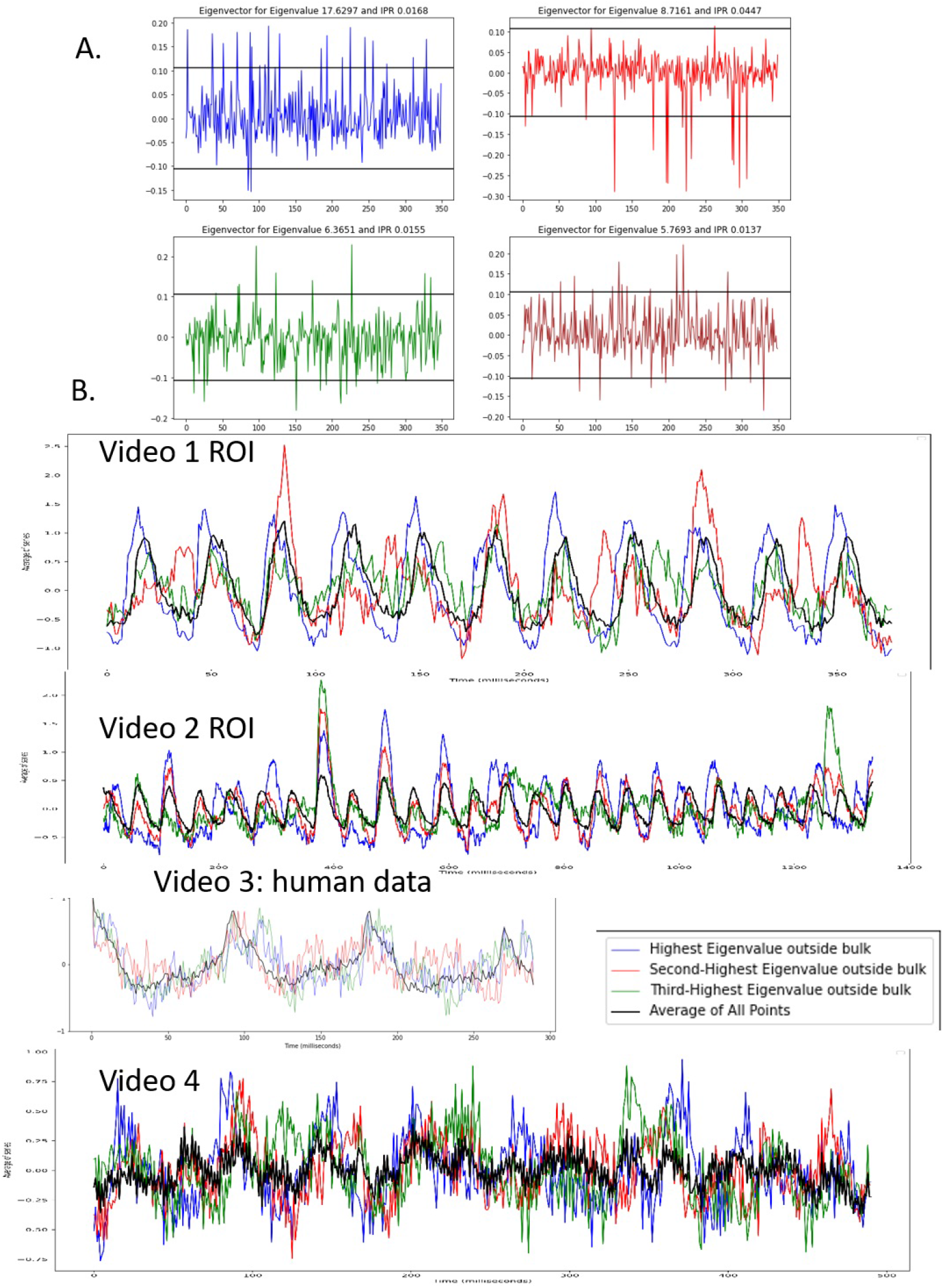
A. Identification of signalling modules in Video 1 i.e. places of localization of the eigenvectors corresponding to high eigenvalues; B: average time series from the signalling modules corresponding for Videos 1-4

We then proceed to investigate the time series associated to the signalling modules. We average over the number of points, to obtain commensurate time series. The time series for Video 1 turns out most remarkable of all. The localization of the top eigenvector identifies a signalling module that predict the AP-induced calcium transient in every beat (in blue). Aside from this remarkable result, the average time series corresponding to other high eigenvalue eigenvectors as well as time series in the other videos demonstrate signals which are heterogeneous in amplitude and frequency, as established in [3].

### 3.2. Common Response

The IPRs corresponding to the top eigenvector are low, indicating that the signal occurs in all the spatial pixels and corresponding to a common response. This is a global signal. For further analysis in order to study local calcium signals, we remove the part corresponding to the common response. To do this we use the matrix **C**_*res*_ defined in equation (10). To verify that the top eigenvector indeed corresponds to common response, we study the correlation coefficient of a least squares regression between *y_max_* as defined in equation (8) and the time series representing the total signal mass at each time point *y_ave_*. We obtain very high correlation coefficients for Video 1 ROI (0.9975), Video 2 ROI (0.9896), and Video 3 (0.9686), and somewhat lower for Video 4 (0.7147). On the other hand taking a the correlation coefficients between *y_ave_* and similar projections onto the eigenvectors corresponding to the eigenvalues in the bulk we obtain a distribution with mean 0 and very low standard deviation for all videos: Video 1 ROI (standard deviation 0.0019), Video 2 ROI (standard deviation 0.0016), Video 3 (standard deviation 0.0044), and Video 4 ROI (standard deviation 0.0072). Thus the top eigenvector indeed correlates very strongly with the total signal mass and indicates common response, while the eigenvectors that are in line with random matrix theory predictions do not correlate with total signal mass at all and indicate noise. As a control, we study the IPRs in Figure 3. The blue circle and the red circle indicate the original signal and the time incremented signal while the blue dot and the red dot indicate the IPRs from the residual matrices. We notice that all the dots are exactly in the middle of their respective circles, except for the eigenvector that was removed, namely eigenvector corresponding to an eigenvalue above λ_*max*_ with a low IPR. Thus this procedure of removing the common response is effective, and does not introduce any other effects.

### 3.3. Correlation Coefficient Structure and Spatial Correlations

We study the distribution of non-diagonal covariance coefficients. In Figure 6, we plot in orange the control covariance coefficients, and observe that they are close to the theoretically predicted Gaussian. Compared to the orange control, the blue plot with the original data is heavily skewed to the right, indicating the presence of correlations. The residual correlation coefficients in green are far closer to the control, reflecting the removed global correlations in Videos 1-3. Yet, they are still far from the Gaussian control, and skewed to the right, showing local correlations. We also notice the correlation coefficients in red for the time incremented series, where the time incrementing removes the time correlations and thus also filters out the AP-induced calcium transients. The time incremented histogram is close to the residual one. This feature is observed for Videos 1, 2, and 3. To check that this is not a particular coincidence of our random sample we study 500 different samples from Video 1 ROI and Video 4 and construct histograms of the square error deviation from the control (Fig 6 B). In Video 1, the square error deviation is large for **C**, and much smaller (an order of magnitude) for **C**_*res*_ and **C′** which are close to each other. This demonstrates that the removed top eigenvector indeed corresponds to global response. In Video 4, however, this effect is less pronounced and the square error deviation from control is commensurate for **C** and **C**_*res*_. We speculate in discussion that this is due to a slower signal propagation, yielding stronger time correlations and weaker space correlations.

**Figure 6.**
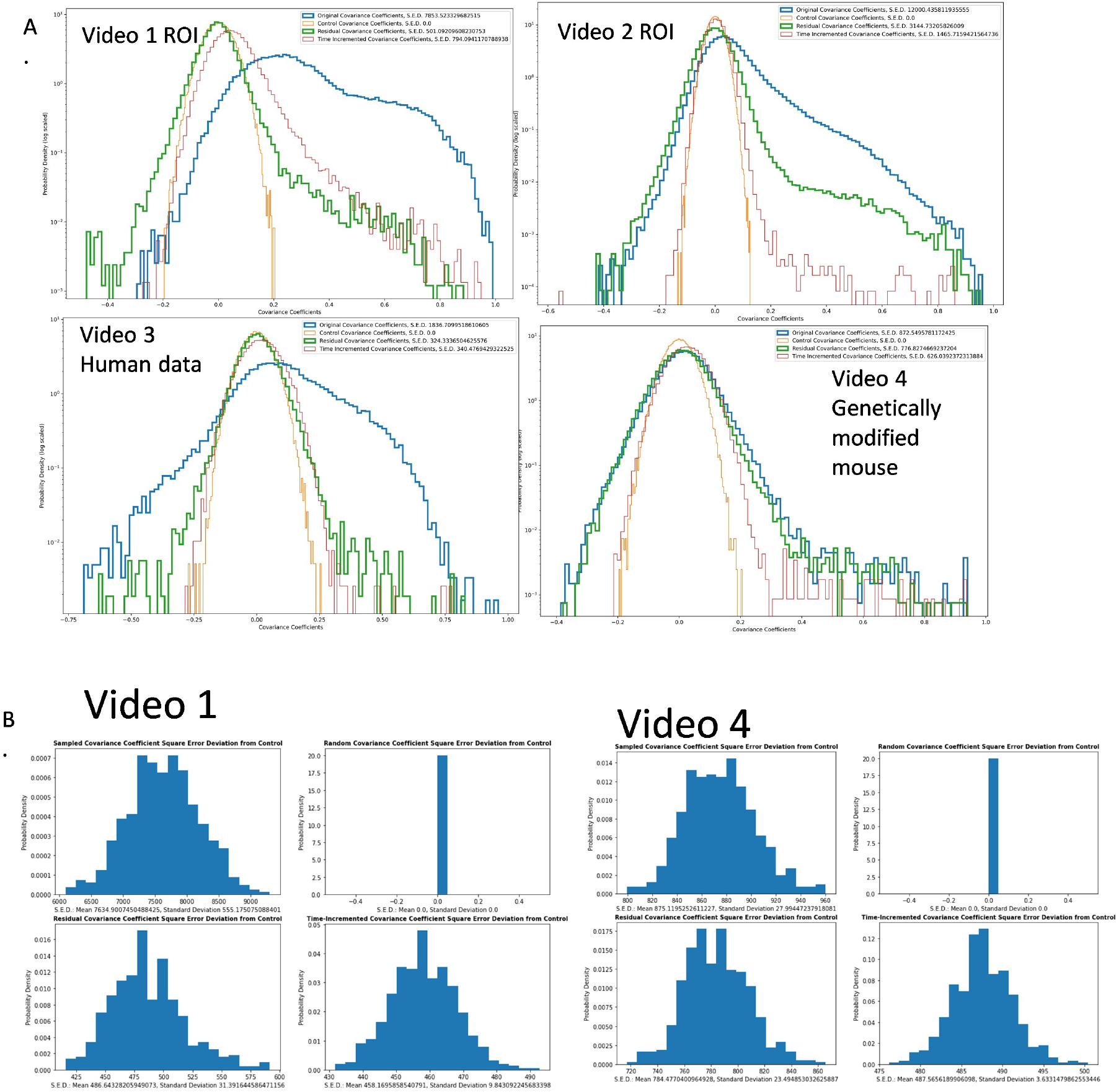
A: Histograms of various covariance coefficients for Videos 1-4. Blue is from the original normalized data sample. Green is from residuals, representing common response to stimulus removed. Red is time-incremented, to remove the common response in a different way. Orange is control from all entries of the original data randomized; B: Total square error of deviation from control (obtained from **C**_*control*_) of all the histograms from A in Videos 1 and 4

Lastly, another important result we can obtain using information on correlation coefficients is the presence of correlations at distances which are higher than cell size. The SAN node cell is known to be roughly spindle shaped at 5-10 *μ*m in diameter and 20-30 *μ*m in length [2]. We sort the correlation coefficients in the upper triangle of **C**_*res*_ according to distance between the pixels and make a separate normalized histogram for pixels with distances 0-20 *μ*m (blue), 20-40 *μ*m (orange), 40-60 *μ* (green), 60-80 *μ*m (red), etc., and we can use far-apart points as control. We see strong deviations from the control in all the distance histograms for Video 1, 2, and 3 (Fig 7 A-C), particularly in the right tails of the distribution. In Fig 7D, we plot the integral of the tail, equivalent to the proportion of correlation coefficients larger than 0.3 for Videos 1, 3, and 4. In Videos 1 and 3, this proportion can be seen to be very high at distances 0 - 20 *μ*m and 20-40 *μ*m. However, the proportions of higher correlations present at distances higher than 40-60 *μ*m and 60-80, visible particularly in the tails plot, indicate intracellular correlations. This proportion is nearly 0 at distances over 80 *μ*m, which can be used as a control. This demonstrates that cells in the central SAN are communicating via calcium. In Video 4, however, the correlations at intercellular distances appear weaker, see Discussion.

**Figure 7.**
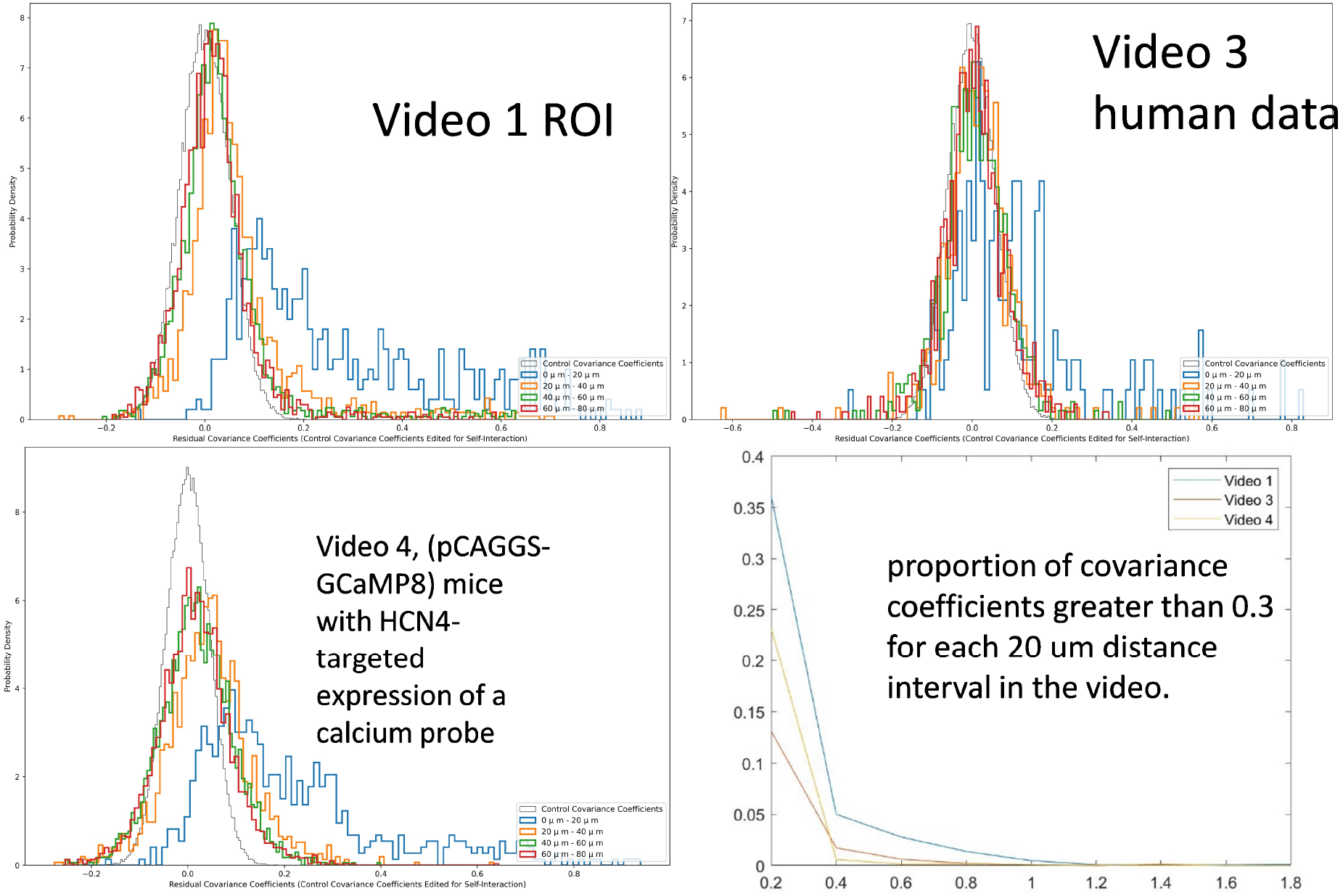
Histograms of covariance coefficients sorted by distance for Videos 1, 2, and 4, blue for 0 to 20 *μ*m, orange for 20-40 *μ*m, green for 40-60*μ*m, red for 60-80 *μ*m, grey for control. Also the integrals of the histogram tails for Videos 1, 3, and 4.

One more check on spatial vs time correlations reveals that space correlations are much more significant than time correlations (Fig 8). The histogram of space correlations is as before constructed from the upper triangle of **C**. To access the time correlations we demean and renormalize the rows instead of the columns, and then construct a *T* × *T* matrix by multiplying the other way. We see that the pink histogram for time correlations is closer to a Gaussian. This means that the time correlations are not too high and our random matrix theory approach is valid. However, as noted before in Video 4 there is a stronger presence of time correlations.

**Figure 8.**
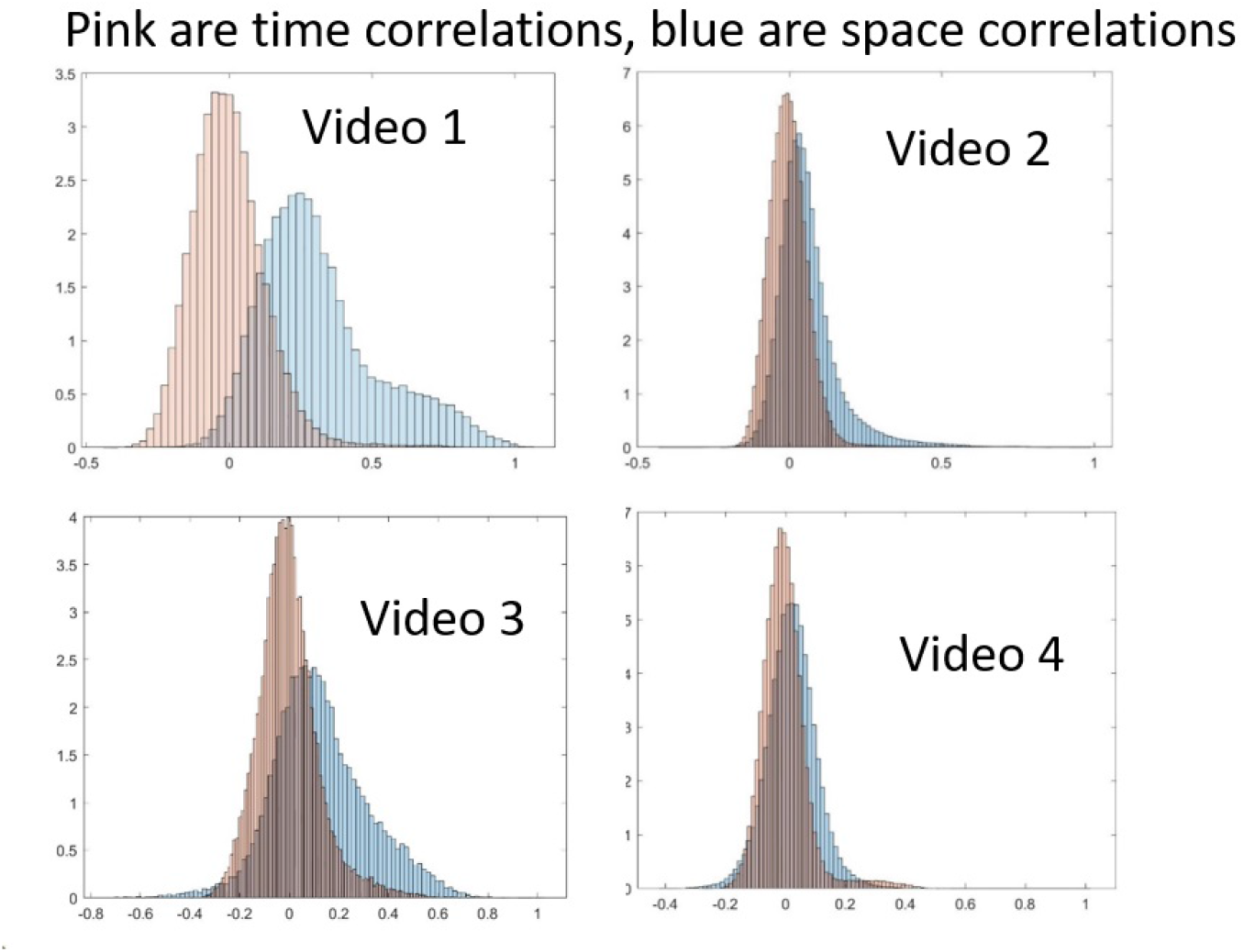
PDF histograms of space correlations (blue) and time correlations (pink) for Videos 1 - 4

## 4. Discussion

In this paper, we have investigated the structure of correlation coefficients between calcium time series at different locations within the SAN using experimental data imaged from intact tissue. The aim is to use random matrix theory to understand properties of calcium signals in SAN and to glean fundamental insights into the workings of SAN. The fundamental question with regard to the SAN is how each action potential transient, which corresponds to a single heart beat, is initiated. It has been shown in [3] that signals at various locations of the SAN are remarkably heterogeneous. Here we take a step toward a mechanistic understanding of the system by demonstrating that the signalling cells interact, thus forming a complex system with heart rate as an emergent phenomenon. How exactly it emerges from the heterogenous signalling units remains unclear, with both the exact biophysical means of cell interaction and the mathematical mechanism of heart rate emergence requiring further research.

Our analysis is based on RMT. We separated the eigenvalues of the signal correlation matrix, and into the parts that are due entirely to noise (those consistent with RMT predictions), those which are due to common response to AP-induced calcium transient (which we can remove for further analysis, e.g. so as to understand the distance-dependence of correlations), and those due to local spatial correlations. We then checked where the eigenvectors are localized indicating points in space where the correlations in the calcium signal changes jointly, thus identifying signalling cell modules. In one of our cases, we the eigenvector localization corresponding to second highest eigenvalue (the highest corresponding to global response)identified a signalling module that initiated the AP-induced calcium transient. This “precursor” module could be a pacemaker module or a signal processing module. It is possible that although we tried to avoid CX43 cells in the ROI, this module is an extension of the “railroad track” CX43 network that carries the excitation. In this case, it is where the central zone first passes the signal to the “rail road track” zone. Further studies could be performed to ascertain how frequently such a “precursor” module is so clearly identified, how it works, and how it is best interpreted.

In our analysis of correlation, we see that Video 4, namely the calcium imaging video from the central area of SAN recorded in preparation from a genetically modified mouse pCAGGS-GCaMP8 with HCN4-targeted expression of Ca2+ probe, has different properties from the other videos. This demonstrates the value of our approach, as these subtle differences in the calcium signals could not be picked up by inspection. We see that the spatial correlations at intercellular distances are smaller than in other videos. We also see that the global response is much less pronounced, as the top eigenvalue is commensurate with the second largest eigenvalue, the top eigenvector is less delocalized, and the correlation coefficient between the signal average and the projection of signals onto the top eigenvector is smaller. These effects could be stemming from differences in the part of the central zone chosen for analysis or from the experimental conditions. We also see that the time correlations in Video 4 are not small compared to the space correlations (Fig 8), which can impact the assumptions of our analysis and could indicate slower signal propagation. This could also be deduced from Fig 2B, as we notice that the number of eigenvalues outside [λ_−_, λ_+_] is much smaller for **C′** than for **C** indicating that the time correlations strongly contribute to this deviation from RMT prediction.

As shown on other systems [13, 8], number variance is a very sensitive parameter. Further studies of the number variance could reveal systematic changes based on various experimental conditions. The number variance could be a characterizing feature of the system and therefore could in the future be useful for diagnostics. However, in using the number variance, we must be cautious of finite size effects, as the number variance for the longest video (Video 2) is the closest to the RMT prediction.

Some of the study limitations are as follows. The analysis was done on only four videos. A future study might conduct this analysis on a large number of videos, and compare the results. It would be interesting to determine how the identified signalling modules of cells interact and create the AP transient. Another limitation is the presence of time correlations which we neglect. We have demonstrated qualitatively that they are small, however, a means of quantitatively establishing how small and how such correlations impact the RMT predictions could be developed in the future. RMT for matrices with dependence in the columns is currently available, cf e.g. [1], and predicts a different distribution of eigenvalues if the dependence is strong enough. This theory can be further developed should there be important statistical applications. Our aim of objective mathematical analysis is only partially achieved, as the identification of ROI was still done by visual inspection. This could be improved, perhaps with machine learning or direct thresholding to identify the central SAN in immunolabeled images. At this point for example it is not clear that the module seen to initiate the signal in Video 1 carries no CX43. Lastly, the histograms for correlation coefficients sorted by distance in Fig 7 have a dependence on the video length, which is why we give the analysis for the three videos whose lengths are commensurate. If there were no time dependence, it is clear that by the Central Limit Theorem the correlation coefficients would scale by 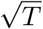. However, since there is time dependence, more sophisticated models of time series must be found or developed to understand the correct scaling. In particular, the coefficients for the longest video (Video 2) exhibit the same pattern as a function of distance, but are smaller by about a factor of 50 in absolute magnitude.

## 5. Conclusions

This paper is a first application of random matrix theory based methods to calcium imaging in the SAN. We borrow and adapt powerful methods that have been used to study correlations various data sets such as finance, EEGs, and calcium imaging in *β*-collective cells. We are able to separate out eigenvalues of the correlation matrix arising from noise, those arising from local correlations, and those indicating a global response. We can furthermore identify signalling modules based on localization properties of eigenvectors and study distance dependence of correlations. This work opens great possibilities for future studies, such as analysis of further data to understand how various interventions might impact the correlation structures of calcium signals. For example, an intervention might change the signalling landscape and the signalling modules might shift. Furthermore, the number variance might be developed into a diagnostic tool, as it is known to change under various experimental conditions.

