## Supplementary material for "Meaningful Local Signaling in Sinoatrial Node Identified by Random Matrix Theory": videos

### Slide 1
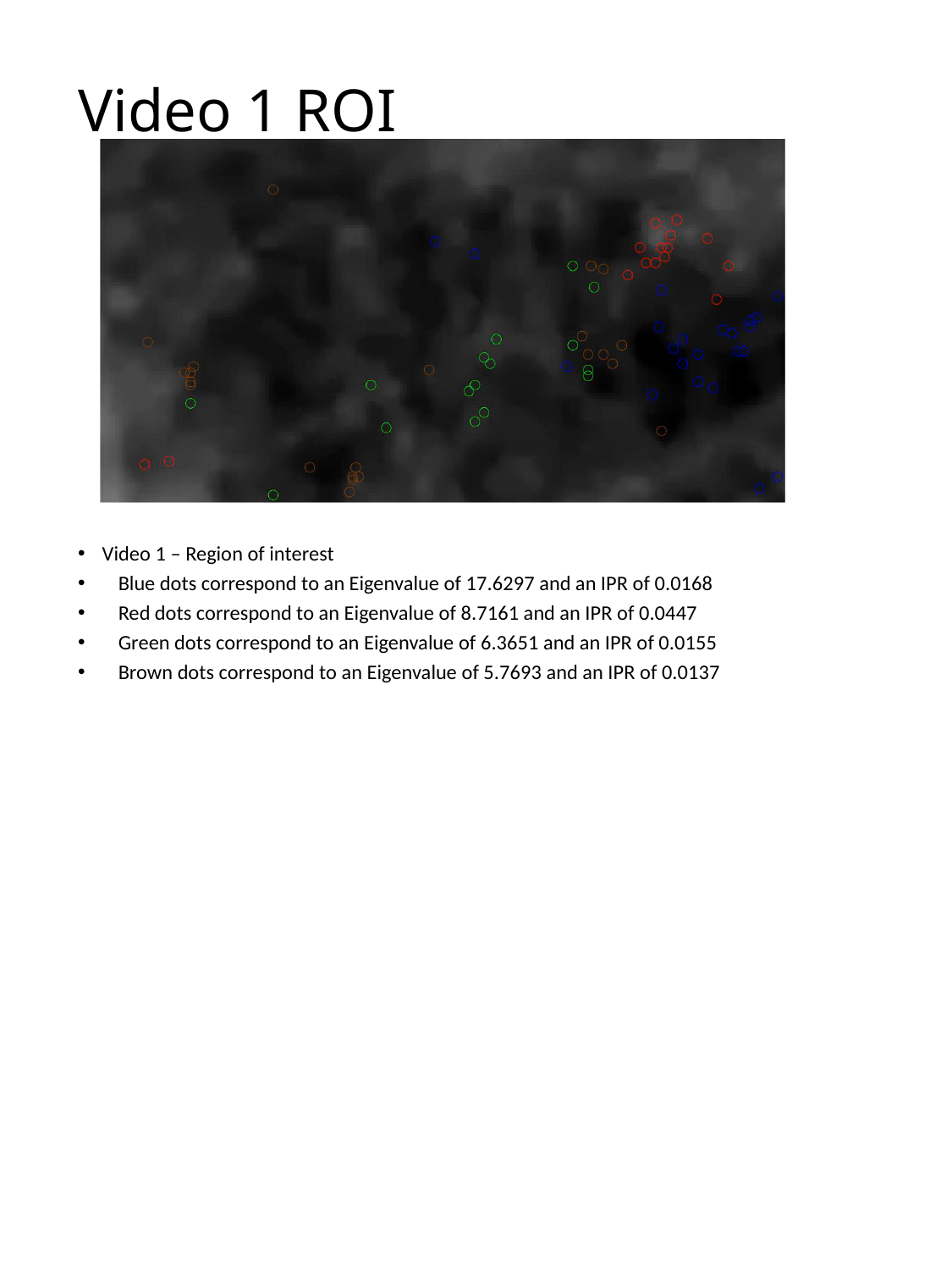

#### Video 1 ROI
Video 1 – Region of interest
Blue dots correspond to an Eigenvalue of 17.6297 and an IPR of 0.0168
Red dots correspond to an Eigenvalue of 8.7161 and an IPR of 0.0447
Green dots correspond to an Eigenvalue of 6.3651 and an IPR of 0.0155
Brown dots correspond to an Eigenvalue of 5.7693 and an IPR of 0.0137

### Slide 2
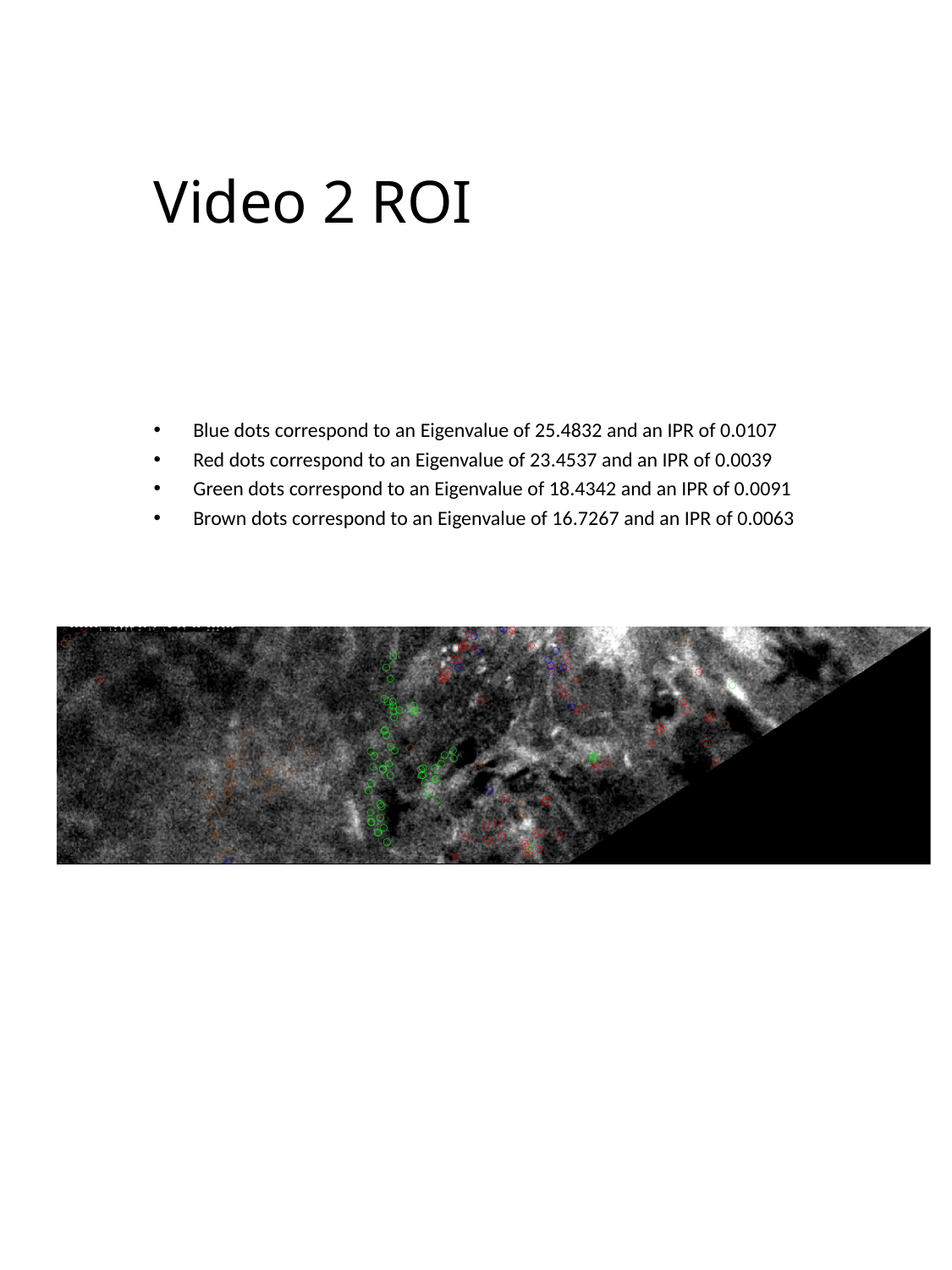

Video 2 ROI
Blue dots correspond to an Eigenvalue of 25.4832 and an IPR of 0.0107
Red dots correspond to an Eigenvalue of 23.4537 and an IPR of 0.0039
Green dots correspond to an Eigenvalue of 18.4342 and an IPR of 0.0091
Brown dots correspond to an Eigenvalue of 16.7267 and an IPR of 0.0063

### Slide 3
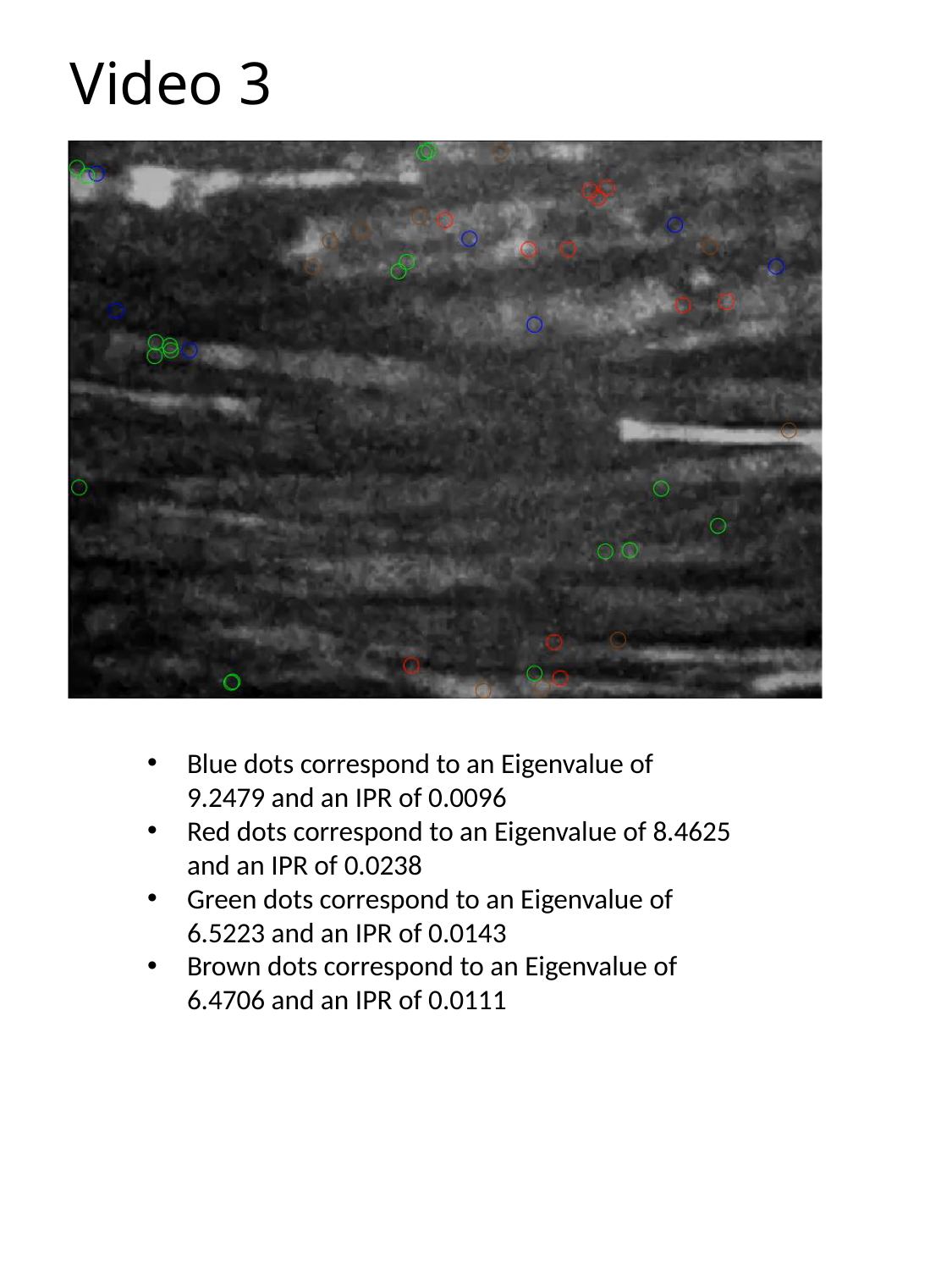

#### Video 3
Blue dots correspond to an Eigenvalue of 9.2479 and an IPR of 0.0096
Red dots correspond to an Eigenvalue of 8.4625 and an IPR of 0.0238
Green dots correspond to an Eigenvalue of 6.5223 and an IPR of 0.0143
Brown dots correspond to an Eigenvalue of 6.4706 and an IPR of 0.0111

### Slide 4
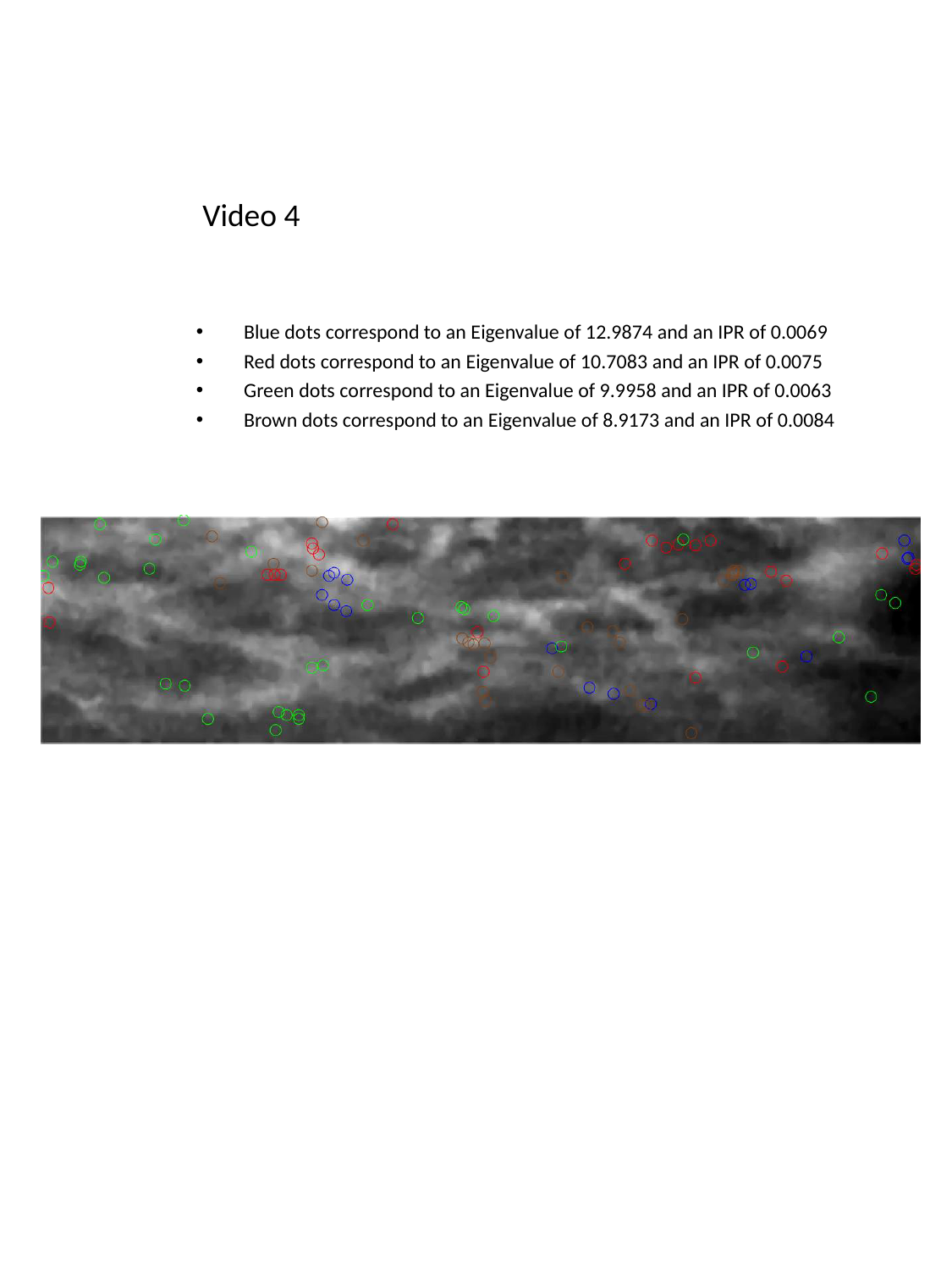

Video 4
Blue dots correspond to an Eigenvalue of 12.9874 and an IPR of 0.0069
Red dots correspond to an Eigenvalue of 10.7083 and an IPR of 0.0075
Green dots correspond to an Eigenvalue of 9.9958 and an IPR of 0.0063
Brown dots correspond to an Eigenvalue of 8.9173 and an IPR of 0.0084

### Slide 5
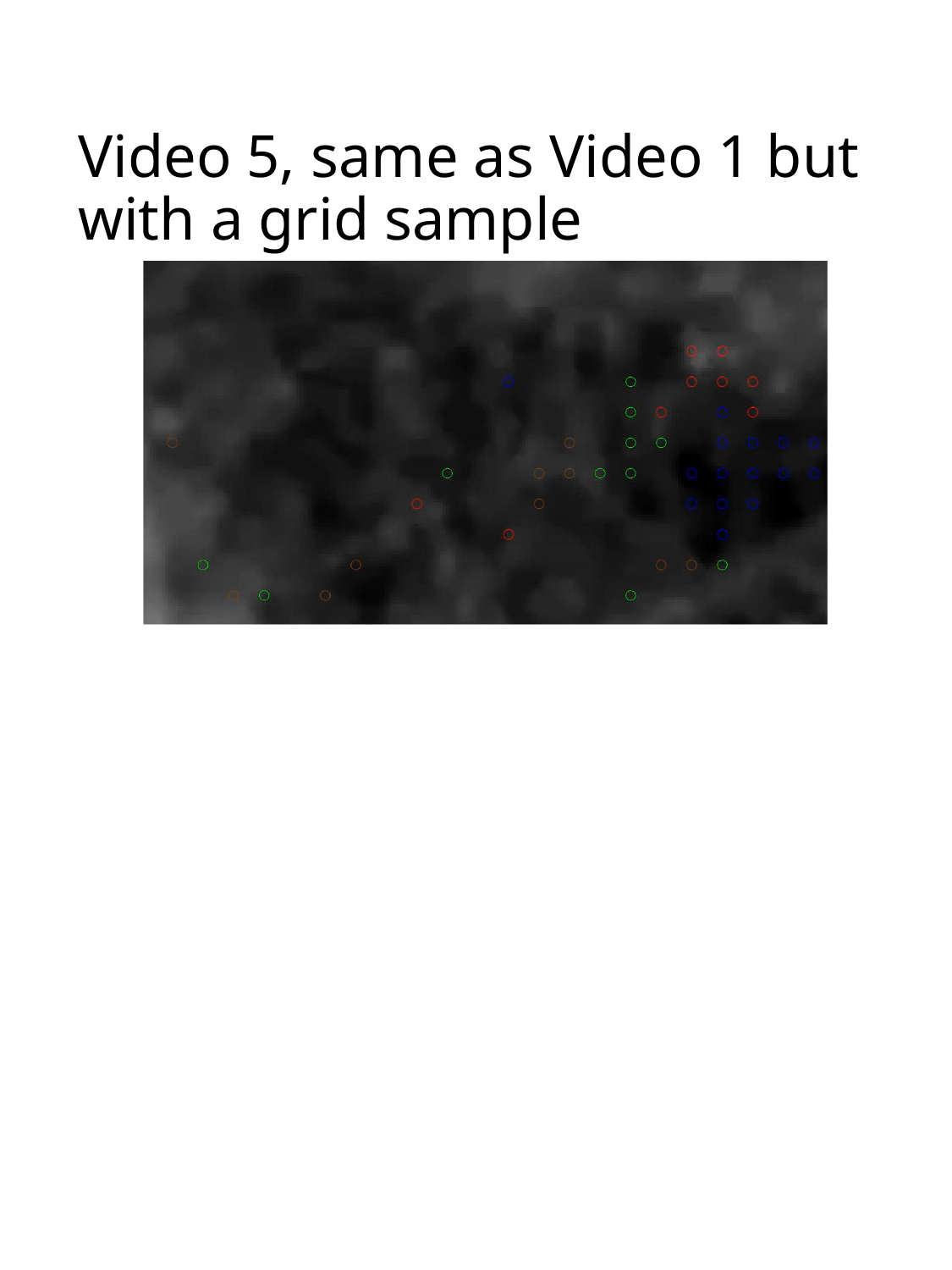

#### Video 5, same as Video 1 but with a grid sample

### Slide 6
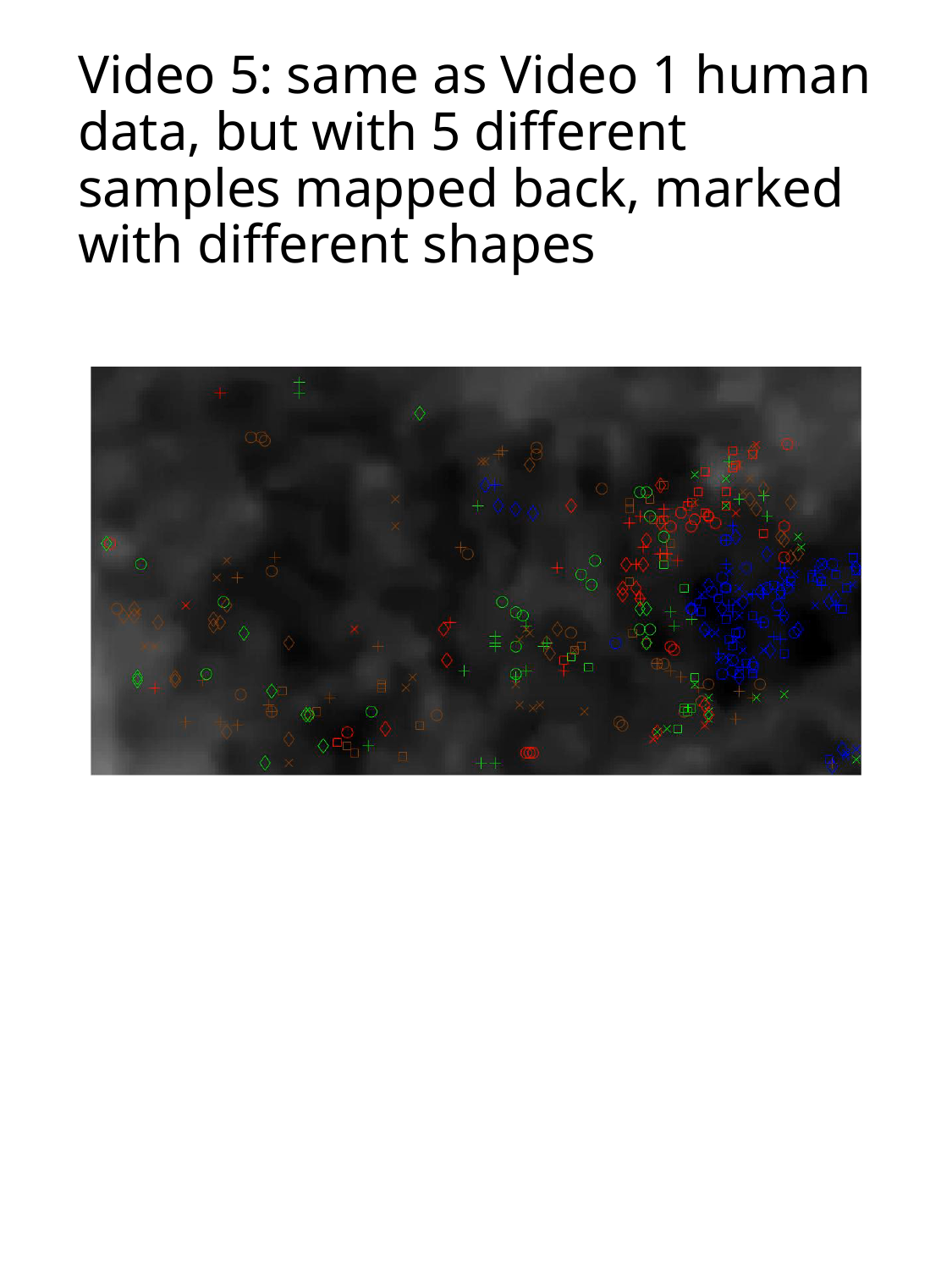

#### Video 5: same as Video 1 human data, but with 5 different samples mapped back, marked with different shapes
